# Increments in Visual Motion Coherence are More Readily Detected than Decrements

**DOI:** 10.1101/2023.01.25.525590

**Authors:** Lai Wei, Autumn O. Mitchell, John H.R. Maunsell

## Abstract

Understanding the circuits that access and read out information in the cerebral cortex to guide behavior remains a challenge for systems-level neuroscience. Recent optogenetic experiments targeting specific cell classes in mouse primary visual cortex (V1) have shown that mice are sensitive to optically-induced increases in V1 spiking, but are relatively insensitive to decreases in neuronal spiking of similar magnitude and time course. This asymmetry suggests that the readout of signals from cortex depends preferentially on increases in spike rate. We investigated whether humans display a similar asymmetry by measuring thresholds for detecting changes in the motion coherence of dynamic random dot stimuli. The middle temporal visual area (MT) has been shown to play an important role in discriminating random dot stimuli, and the responses of its individual neurons to dynamic random dots are well characterized. While both increments and decrements in motion coherence have heterogeneous effects on MT responses, increments cause on average more increases in firing rates. Consistent with this, we found that subjects are far more sensitive to increments of random dot motion coherence than to decrements of coherence. The magnitude of the difference in detectability was largely consistent with the expected difference in effectiveness of coherence increments and decrements in producing increases in MT spike rates. The results add strength to the notion that the circuit mechanisms that read out cortical signals are relatively insensitive to decrements in cortical spiking.

## Introduction

It has long been established that humans and experimental animals can detect electrical activation of small numbers of neurons in their primary visual cortex (V1, reviewed by Histed et al., 2013; Tehovnik & Slocum, 2006). Similarly, it has more recently been shown that mice can detect selective optogenetic activation of their V1 pyramidal neurons (Cone et al., 2020; Cone et al., 2019; Histed & Maunsell, 2014). However, mice have been found to be unexpectedly incapable of detecting decreases in the spike rate of their V1 pyramidal neurons, even when the magnitude of spike rate decreases equaled or exceeded that of readily detected spike rate increases (Cone et al., 2020). This unanticipated asymmetry suggests that the mechanisms decoding activity in the cerebral cortex preferentially monitor increases in spike rate. If so, it would represent an important limitation on what types of cortical information can be accessed to guide behavior.

We explored this issue by determining whether a similar effect can be demonstrated in humans. We identified an opportunity to test the hypothesis of preferential detection of spike rate increases in psychophysical measurements. Here, we describe experiments showing that human subjects are appreciably better at detecting increments in the motion coherence of dynamic random dot stimuli than they are at detecting decrements. This otherwise unanticipated detection asymmetry can be explained by coherence increments producing a much larger increase in the spiking of neurons in the middle temporal visual area (MT) compared to that produced by equivalent coherence decrements (Britten et al., 1992). The results support the hypothesis that increases in cortical spike rates are processed preferentially in guiding behavior.

## Materials and Methods

All experiments were conducted in compliance with a protocol approved by the Institutional Review Board at the University of Chicago and adhered to the tenets of the Declaration of Helsinki. The experimental goals and design were pre-registered before any of the data reported here were collected (Maunsell et al., 2022). The only deviation from the pre-registered design was made after we discovered that dynamic random dots patterns with a 100 ms dot lifetime and a dot density of 2.5 deg^−2^ allowed some subjects to detect changes in the spatial structure of the random dots when coherence changed (see Supplemental Figure 1). The full pre-registered protocol was repeated after adjusting the random dots to have a shorter lifetime (2 frames, 33 ms) and higher density (5 deg^−2^) to obscure the spatial structure of the dots and compel subjects to base their decisions solely on changes in motion coherence.

**Figure 1:**
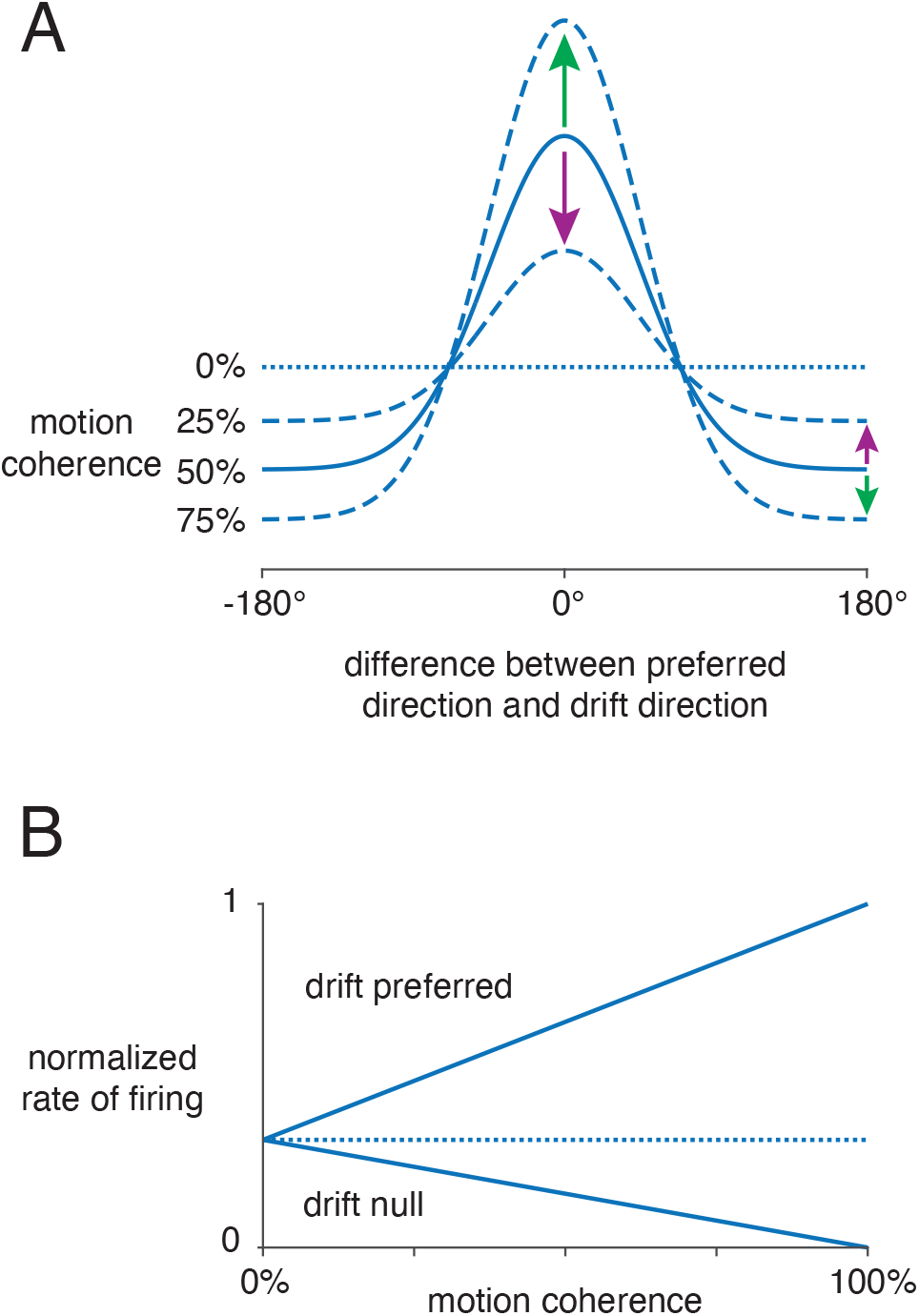
Idealized responses of MT neurons to different random dot motion coherences. **A)** Expected MT average population response to different levels of motion coherence as a function of direction preference relative to dot drift direction. Neurons preferring the drift direction appear at 0°; those preferring the opposite (null) direction appear at ±180°. Four levels of motion coherence are shown. As coherence approaches 0% (dotted line), average responses become unaffected by drift direction because almost all dots move in random directions. Arrows show response changes associated with a step increment in coherence (green, 50% to 75%) or step decrement in coherence (magenta, 50% to 25%). **B)** Effect of varying motion coherence on responses of neurons preferring the drift direction (“drift preferred”) and those preferring the null direction (“drift null”). Responses increase or decrease linearly as motion coherence rises from 0% coherence owing to normalization (see text).

### Choice of Visual Stimuli

We measured the ability of subjects to detect changes in the motion coherence of dynamic random dots. We used random dots because the primate MT has been shown to play a central role in discriminating random dot motion. Inactivation of MT substantially impairs dot motion discrimination (Newsome & Pare, 1988), and selective activation of MT subpopulations preferring a particular direction of motion can enhance the perception of that direction (Salzman et al., 1992). The performance of monkeys in discriminating directions of motion of dynamic random dots closely matches the performance of individual MT neurons in discriminating directions of motion (Britten et al., 1992).

Importantly, MT neurons also have distinctive response properties that allow us to predict the relative magnitude of their spike rate changes in response to changes in the motion coherence of dynamic random dots. When motion coherence for a given direction of dot drift is incremented, neurons preferring that direction will increase their rate of firing, while neurons preferring the opposite (null) direction will typically decrease their rate of firing (Figure 1A, green arrows). In most cases, an increment in coherence will increase the spike rate of neurons preferring the drift direction by far more than it decreases the spike rate of neurons preferring the null direction. Conversely, a decrement in coherence will increase the spike rate of neurons preferring the null direction by far less than it decreases the spike rate of neurons preferring the drift direction (Figure 1A, magenta arrows). This difference was described by Britten and his colleagues (1993), who found that only about half their recorded neurons had a statistically significant suppression of firing to null direction motion. They additionally noted that the firing rates of most MT neurons vary linearly with motion coherence, with neurons preferring the drift direction increasing their rate of firing about 3.5 times faster than neurons preferring the null direction decrease their rate of firing (0.39 versus −0.11 spikes/s/percent-coherence).

This linear relationship between spike rate and motion coherence follows from the robust normalization that MT neurons show when more than one stimulus appears in their receptive field (Ni et al., 2012; Rust et al., 2006). If each dot in a dynamic random dot display is viewed as an individual stimulus, then the divisive normalization model (Heeger, 1992) predicts the response will be:

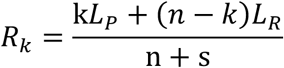

where n is the number of dots, k is the number moving in the preferred direction, s is a semi-saturation constant, L_P_ corresponds to the strength of response to preferred direction (coherent) motion and *L_R_* corresponds to the strength of response to random directions of motion (0% coherent). *L_R_* is different from the response to motion in null direction (*L_N_*), and takes a value between *L_P_* and *L_N_*. Transferring one stimulus dot from the 0% coherent pool to the coherent pool increases the numerator by a fixed amount (*L_P_* − *L_R_*) regardless of the coherence, resulting in a linear increase in *R_k_* with coherence. For coherent motion in the null direction, increasing coherence will linearly reduce response because (*L_N_* − *L*_R_) will be negative. Britten and his colleagues (1993) observed a strong neuron-by-neuron correlation between response to 0% coherent motion and the response to 100% coherent motion averaged over each of eight directions, which is expected from response normalization.

A greater rate of change in spike rate for motion in the preferred relative to null direction arises because the response to 0% coherent motion is typically closer to the null direction response than to the preferred direction response (Figure 1B). Normalization dictates that a neuron’s response to 0% coherent motion will be the average of its responses across the full range of directions. This average will depend on the width of the neuron’s direction tuning. When direction tuning is extremely sharp, the average response across all directions approaches the null direction response. Conversely, as direction tuning is extremely broad, the average response across all directions approaches the preferred direction response. If we assume MT direction tuning follows a Gaussian function and take the preferred and null direction responses to be 1 and 0, the response to 0% coherent motion will be:

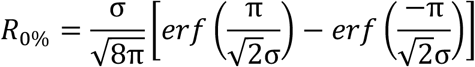

which is the average response over the range ±180°. As expected, *R*_0%_ depends on the width of the tuning curve (σ). Albright (1984) measured the direction tuning of monkey MT neurons using dynamic random dots, and found an average tuning width corresponding to σ = 36°. Britten and Newsome (1998) similarly found an average tuning width of σ = 39°. These values yield *R*_0%_ equal to 0.25 and 0.27 times the response to 100% coherent motion in the preferred direction, suggesting a roughly 3:1 difference in the change in rate of firing for neurons preferring the drift direction and the opposite direction when coherence changes. This ratio is slightly less than the values reported by Britten and colleagues (1993), who found the average MT response to 0% coherence motion to be 0.20 times the response to 100% coherent preferred direction motion and a 3.5:1 difference in change in rate of firing to preferred versus null direction dot motion.

These calculations are based entirely on measurements of single unit responses in macaque monkeys because recordings from single units in human MT are not available. However, the direction selectivity of MT neurons is highly conserved across primate species (Allman et al., 1985; Lui & Rosa, 2015; Rosa & Elston, 1998), and fMRI recordings have shown that human MT neurons have similar direction selectivity (Tootell, Reppas, Dale, et al., 1995) and contrast sensitivity (Tootell, Reppas, Kwong, et al., 1995) to that in non-human primates.

The asymmetric changes in MT spike rates for preferred and null direction motion provide a test for the idea that detection of stimulus changes depends preferentially on increasing spike rates. When motion coherence is incremented, MT neurons preferring the drift direction will have the largest increase in firing rate (Figure 1A, green arrow at 0°). However, when motion coherence is decremented, the largest increase in firing rate will occur in neurons that prefer motion opposite to the drift direction, and that increase in firing rate will be much smaller (Figure 1A, magenta arrow at 180°). Importantly, the linear relationship between MT spikes rates and coherence ensures that this relationship holds regardless of the size or starting point of the coherence change (Figure 1B). Evoking the same increase in spiking with a decrement in coherence will require a much larger coherence decrement compared to a coherence increment. Because we anticipate that perceptual thresholds will correspond to a given change in MT neuron firing rates (Britten et al., 1992), we expect them to show a corresponding asymmetry. Alternatively, if increases and decreases in spike rates are equally detectable, increments and decrements in coherence should have similar thresholds.

### Experimental Design

Subjects were recruited from the university community and were naive about the goals of the experiments. The primary data were collected from 5 subjects (22-27 years old, 2 male) with normal or corrected to normal vision. Two other subjects were unable to achieve stable performance over the course of an initial one-hour training session and were dropped from the study. Subjects viewed a calibrated computer display (1920 × 1080 pixels, 60 Hz frame rate) and used a chin rest to maintain a viewing distance of 57 cm. Each trial began with the appearance of a central fixation spot, on which the subject held their gaze throughout the trial. The subject pressed a key to signal when they had fixated and were ready, causing two patches of dynamic random dots to appear on either side of the fixation spot (10° diameter centered on azimuth ±10°, elevation 0°, 5 dots/deg^2^, dot diameter 0.1°, dot life two frames, 33 ms, speed 16 deg/s, white dots on a mid-level gray background). The parameters were selected to match MT receptive field size (Albright & Desimone, 1987) and speed preferences (Maunsell & Van Essen, 1983). Except as noted, both patches initially drifted upward with 50% coherence. After 1 s, the coherence of a randomly selected patch changed for 250 ms, after which both patches disappeared. Subjects had to signal which patch had changed coherence using the left or right arrow key on a keyboard. Subjects were under no time pressure to respond and were encouraged to take frequent breaks.

Thresholds for detecting coherence increments and decrements were determined using a 3-down, 1-up staircase. The initial coherence change was selected randomly, and initial staircase steps were 8% coherence. The staircase step size was halved after each six reversals, to a minimum of 1%. Threshold was taken as the coherence change on the 100^th^ trial. Increment and decrement thresholds were measured on alternating days so that subjects were fully aware of the change that they were looking for on each trial. In the primary data set, the five subjects each provided five threshold measurements during each of six sessions (15 increment thresholds, 15 decrement thresholds).

## Results

Each of the five subjects had a markedly larger average threshold for detecting coherence decrements than for detecting increments (Figure 2). Across all subjects, the average decrement threshold was 39.3% (1.0% SEM), while the average increment threshold was 20.3% (0.5% SEM; average ratio 1.94, range 1.75-2.19). Larger decrement thresholds are consistent with spike increases in MT being more readily detected than spike rate decreases (Figure 1).

**Figure 2:**
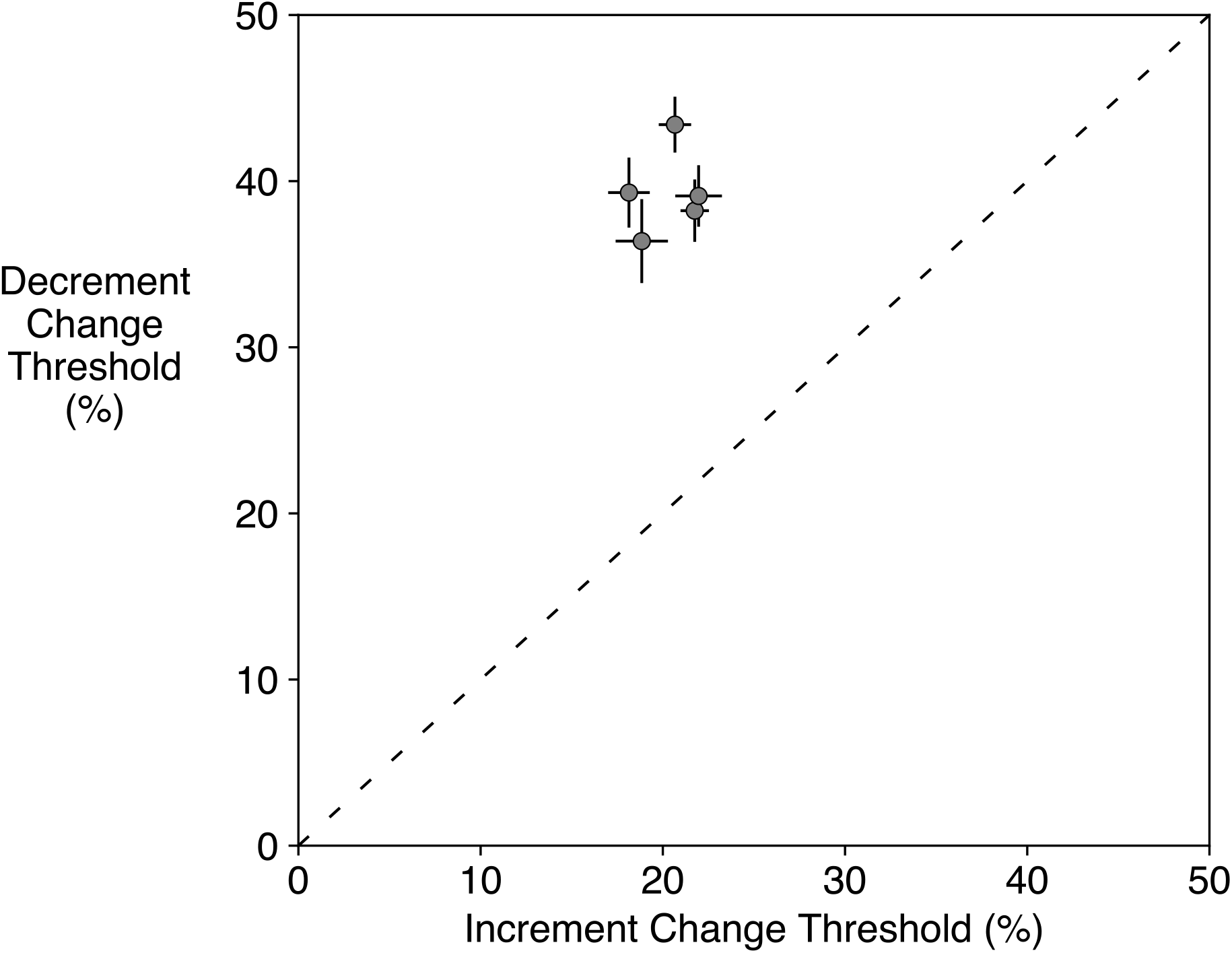
Average thresholds for detecting steps of motion coherence increments and decrements. Each point shows the average thresholds for one of the five subjects (±1 SEM). Thresholds were consistently larger for coherence decrements. The aggregate mean increment threshold was 20.3%, while the aggregate mean decrement threshold was 39.3% (ratio 1.94, effect size 2.9).

The thresholds shown in Figure 2 were measured as steps up or down from 50% coherence. Thus, increment detections involved ranges of motion coherence that did not overlap with the ranges of motion coherence used for decrement detections. It is possible that different coherence ranges used could affect the thresholds. For example, when the initial coherence is 100%, detecting a decrement becomes the trivial task of spotting a single dot moving in a different direction. To address this concern, we ran two of the subjects in additional sessions in which we measured five increment thresholds starting from a coherence that was equal to 50% coherence minus their average decrement threshold, thereby ensuring that the coherences used for these increment thresholds were entirely contained within the range of coherences used for the corresponding decrement thresholds. In neither case did overlapping the coherence ranges for increments and decrements appreciably reduce the ratio of the thresholds (Subject 1: decrement threshold from 50%, 43.4%; increment threshold from 50% coherence, 20.7%; increment threshold from 6.4%, 21.0%; Subject 2: decrement threshold from 50%, 36.3%; increment threshold from 50% coherence, 18.9%; increment threshold from 13.7%, 13.8%). This consistency in thresholds shows that the threshold difference does not depend on differences in the range of coherences the increment and decrement steps cover.

We also considered the possibility that subjects might improve over the course of the sessions, with faster improvement on detecting increments than on detecting decrements. Were that the case, we might see a difference in the two thresholds over 15 sessions even if asymptotic thresholds were identical. To examine this, we fit exponentials to the across-subject averages for the 15 threshold measurements, separately for increment and decrement thresholds. The exponents for the increment and decrement fits were 0.8 and 1.0, indicating little improvement for either threshold past the first few measurements. Consistent with this, the asymptotes for the two fits (i.e., thresholds extrapolated to infinite training) closely matched the values reported above (increments 19.5% and decrements 38.5%; ratio 1.97).

## Discussion

We found that thresholds for detecting decrements in the coherence of dynamic random dots are substantially greater than thresholds for detecting increments of coherence. Ideal observer models of dynamic random dot thresholds have focused primarily on direction discrimination (Barlow & Tripathy, 1997; Watamaniuk, 1993), but an ideal observer would be expected to have equal thresholds for coherence increments and decrements because they are simply time-reversed instances of each other. Given the expected responses of MT neurons to these stimuli (Materials and Methods), this result supports findings from optogenetic stimulation experiments that show mice detect increases in neuronal spiking in visual cortex preferentially over decreases in spiking (Cone et al., 2020). Together, they suggest that insensitivity to decreases in cerebral cortex spiking applies to all mammals.

We observed relatively little scatter in the ratio of measured coherence thresholds (Figure 2). However, the average ratio between coherence decrement and increment thresholds (1.9 fold) was appreciably smaller than that predicted based on a Gaussian fit to the responses of MT neurons to different directions of dynamic random dot stimuli (3 fold: Albright, 1984, Britten & Newsome 1998, see Materials and Methods). One explanation for this difference is that the readout of neuronal responses in MT might necessarily pool responses from neurons preferring a range of directions. The predicted ratio of 3 assumes perfect access to only those neurons that preferred the dot drift direction and its opposite (arrows in Figure 1A). If responses are instead pooled across ranges around these direction preferences, the ratio of spike rate changes will be smaller. Assuming a uniform weighting of spikes across a range of direction preferences for Gaussian tuning curves with σ = 37.5° (average of Albright 1984 and Britten & Newsome 1998), the ratio of changes in spike rates around the preferred and null directions drops to the observed average psychophysical ratio for a pooling range of ±50.2° around these directions. This is a broad range, but approximates the range of direction preferences over which neurons increase their rate of firing in response to motion coherence increments (±40.2° around preferred drift direction when σ = 37.5°; curves above 0% coherence response in Figure 1A).

Several other factors might also contribute to a difference between expected and observed threshold ratios. Although MT plays an important role in the assessment of dynamic random dot motion (Britten et al., 1992; Newsome & Pare, 1988; Salzman et al., 1992), motion perception survives ablation of MT (Newsome & Pare, 1988; Pasternak & Merigan, 1994). If direction tuning in other cortical areas supporting motion perception were broader than that in MT, their contribution might produce more similar coherence increment and decrement thresholds (see Materials and Methods). Additionally, noise downstream from perceptual processing (e.g., lapses) would also reduce the differences between the thresholds. It is also possible that the response properties of human MT neurons differ from those of the macaque MT neurons in which the estimates were based. Finally, some subjects spontaneously reported that they found decrement detection more difficult, and it is possible they attended more to the task when working with those stimuli. These potential factors are not mutually exclusive, and all might contribute to a smaller threshold ratio than predicted based solely on the direction tuning curves of MT neurons. Given uncertainties about factors like these, the lack of a precise quantitative match between MT tuning and behavioral thresholds is unsurprising. Ultimately, the highly significant threshold difference supports the observations from mouse experiments indicating that decrements in cerebral cortical spiking are appreciably less well detected than increases in spiking.

It is possible that preferential sensitivity to spike rate increases is widespread in the brain. Stimulation studies in rodents (Cone et al., 2020; Dalgleish et al., 2020; Houweling & Brecht, 2008; Huber et al., 2008) and monkeys (DeYoe et al., 1989; Murphey & Maunsell, 2007, 2008; Ni & Maunsell, 2010) have shown that animals can respond to the excitation of small numbers of neurons in all regions of the cerebral cortex, as well as a broad spectrum of monkey subcortical areas (Doty, 1961; Doty, 1965; Nielson et al., 1962). Such a preference might be driven by energetic considerations. The energy costs of neuronal spiking are extraordinarily high (Attwell & Laughlin, 2001), limiting the average sustained rate of spiking for cortical neurons to less than one spike a second (Lennie, 2003). Recurrent neural networks trained to do stimulus detection rely on activity increments when neural activity has appreciable costs (Cone et al., 2020). Preferentially processing spike rate increases might also confer computational robustness. Examination of the performance in artificial spiking networks has shown that circuits that are insensitive to spike rate decreases can provide highly reliable decoding (Calaim et al., 2022). Similarly, decoders that use only increases in spike rate can do an excellent job of detecting and decoding stimulus and motor events (Sadras et al., 2019). More broadly, the many detection models that signal when a leaky integrator of spike counts reaches a threshold level of activation (e.g., Cook & Maunsell, 2002; Hanes & Schall, 1996) typically will not detect decreases in spike rate.

These observations notwithstanding, there is little reason to expect insensitivity to spike rate decreases is universal in the brain. For example, some Purkinje cells in the cerebellum maintain an average high rate of simple spike firing that provides a continuous time-varying signal of motor errors (Popa et al., 2016). It is possible that the decoding of increases and decreases of those rates are equally important for motor precision, and the same might apply for all neurons that modulate high sustained rates of firing. Similarly, decreases in firing rates in the structures like the nucleus accumbens (Krause et al., 2010) and substantia nigra (Hikosaka, 2007) play a permissive role in initiating behavioral responses.

The previous results from mouse optogenetics (Cone et al., 2020) and the current results from human psychophysics suggest that spike rate decreases in the visual cortex might not be accessible for behavioral detection. However, Luis-Islas and colleagues (2022) recently showed that mice expressing ChR2 under the control of the vesicular GABA transporter (VGAT) could learn to detect optical stimulation of their prefrontal cortex, despite the fact that the primary effect of that stimulation was to reduce neuronal firing rates. Although the mice learned to detect stimulation of VGAT stimulation, learning rates were significantly slower than those for stimulation of pyramidal neurons (ChR2 expression under control of mouse thymus antigen 1 promoter, Thy-1). Nevertheless, this leaves the question of whether spike rate decreases are undetected or only less well detected. Several factors might account for the difference between the studies. Cone and colleagues (2020) reported that mice failed to detect activation of parvalbumin-expressing (PV) or somatostatin-expressing (SST) cortical neurons. VGAT exists in vasopressin-expression (VIP) neurons in addition to PV and SST cells. Mice can reliably report the activation of VIP neurons (Cone et al., 2019), presumably because VIP neuron activation increases pyramidal cell spiking (Pi et al., 2013). Thus, co-activation of PV, SST and VIP neurons in VGAT mice might have increased the spiking of at least some pyramidal neurons. Also, the VGAT study used relatively strong optogenetic stimulation (1800 μJ over 1000 ms compared with 125 μJ over 500 ms in Cone et al., 2020). When the intense and sustained suppression ended, it caused a rebound in spiking that might have contributed to the perceptual reports. While both studies point to preferential behavioral detection of spike rate increases, more studies will be needed to establish with precision the extent to which increases in cortical spike rates are preferentially detected over decreases in spike rates. Progress on this question might be made through direct neurophysiological recordings and perturbations of MT neurons in monkeys while they perform coherence change detection.

## Acknowledgments

Supported by the Neuroscience Institute of the University of Chicago. We thank Marlene R. Cohen for insightful discussions regarding the interpretation of these results, and Chery Cherian, Jackson J. Cone and Supriya Ghosh for helpful comments on drafts of the manuscript. LW was supported by a Dean’s International Student Fellowship from the Biological Sciences Division at the University of Chicago.

## Supplementary Material

**Supplemental Figure 1:**
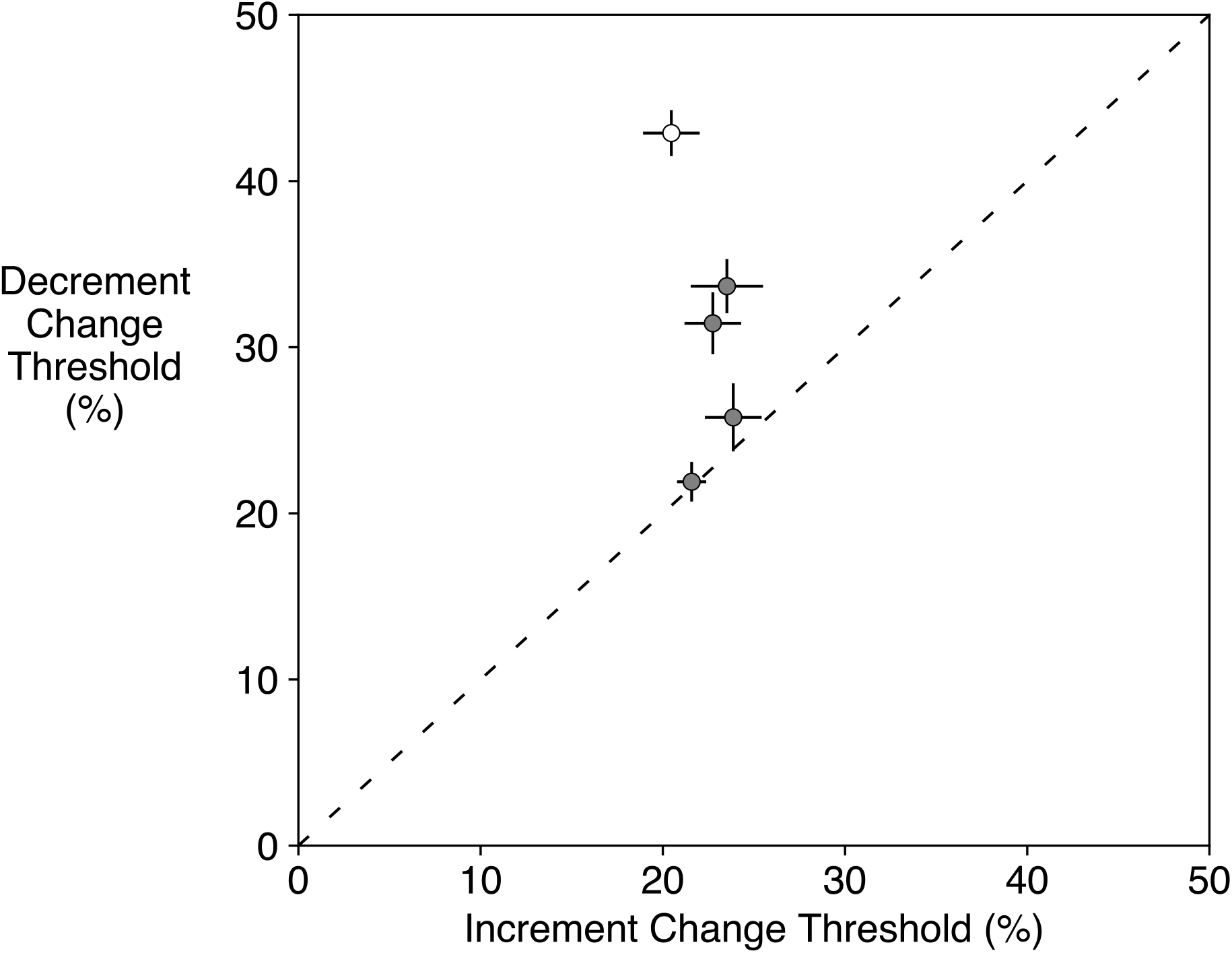
Average thresholds for detecting steps of motion coherence increments and decrements using dynamic random dots with 100 ms lifetime and 2.5 dot/deg^2^. Each point shows the average thresholds for one of five subjects (±1 SEM). The dot lifetime and density allowed some subjects to detect changes in the spatial structure of the random dots when coherence changed and achieve similar change detection thresholds regardless of the sign of the change. The data reported in the main text were collected from 4 of the same 5 subjects (filled symbols) after adjusting the random dots to have a shorter lifetime and higher density to remove the spurious cues. The remaining subject (open symbol) was unable to achieve stable coherence decrement thresholds when those cues were removed and was replaced.

## Literature Cited

Albright, T. D. (1984, Dec). Direction and orientation selectivity of neurons in visual area MT of the macaque. J Neurophysiol, 52(6), 1106–1130. https://doi.org/10.1152/jn.1984.52.6.1106

Albright, T. D., & Desimone, R. (1987). Local precision of visuotopic organization in the middle temporal area (MT) of the macaque. Experimental Brain Research, 65, 582–592.

Allman, J. M., Miezin, F. M., & McGuinness, E. (1985). Direction- and velocity-specific responses from beyond the classical receptive field in the middle temporal visual area (MT). Perception, 14, 105–126.

Attwell, D., & Laughlin, S. (2001). An energy budget for signaling in the grey matter of the brain. Journal of Cerebral Blood Flow Metabolism, 21, 1133–1145.

Barlow, H. B., & Tripathy, S. P. (1997). Correspondence noise and signal pooling in the detection of coherent visual motion. Journal of Neuroscience, 17, 7954–7966.

Britten, K. H., & Newsome, W. T. (1998). Tuning bandwidths for near-threshold stimuli in area MT. Journal of Neurophysiology, 80, 762–770.

Britten, K. H., Shadlen, M. N., Newsome, W. T., & Movshon, J. A. (1992). The analysis of visual motion: A comparison of neuronal and psychophysiology performance. Journal of Neuroscience, 12, 4745–4765.

Britten, K. H., Shadlen, M. N., Newsome, W. T., & Movshon, J. A. (1993). Responses of neurons in macaque MT to stochastic motion signals. Visual Neuroscience, 10, 1157–1169.

Calaim, N., Dehmelt, F. A., Goncalves, P. J., & Machens, C. K. (2022, May 30). The geometry of robustness in spiking neural networks. eLife, 11. https://doi.org/10.7554/eLife.73276

Cone, J. J., Bade, M. L., Masse, N. Y., Page, E. A., Freedman, D. J., & Maunsell, J. H. R. (2020, Oct 7). Mice Preferentially Use Increases in Cerebral Cortex Spiking to Detect Changes in Visual Stimuli. J Neurosci, 40(41), 7902–7920. https://doi.org/10.1523/JNEUROSCI.1124-20.2020

Cone, J. J., Scantlen, M. D., Histed, M. H., & Maunsell, J. H. R. (2019, Jan-Feb). Different Inhibitory Interneuron Cell Classes Make Distinct Contributions to Visual Contrast Perception. eNeuro, 6(1). https://doi.org/10.1523/ENEURO.0337-18.2019

Cook, E. P., & Maunsell, J. H. R. (2002). Dynamics of neuronal responses in macaque MT and VIP during motion detection. Nature Neuroscience, 5, 985–994.

Dalgleish, H. W., Russell, L. E., Packer, A. M., Roth, A., Gauld, O. M., Greenstreet, F., Thompson, E. J., & Hausser, M. (2020, Oct 26). How many neurons are sufficient for perception of cortical activity? eLife, 9. https://doi.org/10.7554/eLife.58889

DeYoe, E. A., Lewine, J. D., & Doty, R. W. (1989). Optimal stimuli for detection of intracortical currents applied to striate cortex of awake macaque monkeys. Proceedings of the Annual International Conference of IEEE Engineering in Medicine and Biology Society, 11, 934–936.

Doty, R. W. (1961, Jul 28). The role of subcortical structures in conditioned reflexes. Ann N Y Acad Sci, 92, 939–945. https://doi.org/10.1111/j.1749-6632.1961.tb40967.x

Doty, R. W. (1965). Conditioned reflexes elicited by electrical stimulation of the brain in macaques. Journal of Neurophysiology, 28, 623–640.

Hanes, D. P., & Schall, J. D. (1996). Neural control of voluntary movement initiation. Science, 274, 427–430.

Heeger, D. J. (1992). Normalization of cell responses in cat striate cortex. Visual Neuroscience, 9, 181–197.

Hikosaka, O. (2007). GABAergic output of the basal ganglia. Prog Brain Res, 160, 209–226. https://doi.org/10.1016/S0079-6123(06)60012-5

Histed, M. H., & Maunsell, J. H. R. (2014, Jan 07). Cortical neural populations can guide behavior by integrating inputs linearly, independent of synchrony. Proceedings of the National Academy of Science USA, 111(1), E178–187. https://doi.org/10.1073/pnas.1318750111

Histed, M. H., Ni, A. M., & Maunsell, J. H. R. (2013, Apr). Insights into cortical mechanisms of behavior from microstimulation experiments. Prog Neurobiol, 103, 115–130. https://doi.org/10.1016/j.pneurobio.2012.01.006

Houweling, A., & Brecht, M. (2008, Jan 3). Behavioural report of single neuron stimulation in somatosensory cortex. Nature, 451(7174), 65–68. http://www.ncbi.nlm.nih.gov/entrez/query.fcgi?cmd=Retrieve&db=PubMed&dopt=Citation&list_uids=18094684

Huber, D., Petreanu, L., Ghitani, N., Ranade, S., Hromadka, T., Mainen, Z., & Svoboda, K. (2008, Jan 3). Sparse optical microstimulation in barrel cortex drives learned behaviour in freely moving mice. Nature, 451(7174), 61–64. http://www.ncbi.nlm.nih.gov/entrez/query.fcgi?cmd=Retrieve&db=PubMed&dopt=Citation&list_uids=18094685

Krause, M., German, P. W., Taha, S. A., & Fields, H. L. (2010, Mar 31). A pause in nucleus accumbens neuron firing is required to initiate and maintain feeding. J Neurosci, 30(13), 4746–4756. https://doi.org/10.1523/JNEUROSCI.0197-10.2010

Lennie, P. (2003). The cost of cortical computation. Current Biology, 13, 493–497.

Lui, L. L., & Rosa, M. G. (2015, Apr). Structure and function of the middle temporal visual area (MT) in the marmoset: Comparisons with the macaque monkey. Neurosci Res, 93, 62–71. https://doi.org/10.1016/j.neures.2014.09.012

Luis-Islas, J., Luna, M., Floran, B., & Gutierrez, R. (2022, May-Jun). Optoception: Perception of Optogenetic Brain Perturbations. eNeuro, 9(3). https://doi.org/10.1523/ENEURO.0216-22.2022

Maunsell, J. H. R., Mitchell, A., & Wei, L. (2022). Detection of neuronal firing rate increments and decrements. https://osf.io/3wu7f

Maunsell, J. H. R., & Van Essen, D. C. (1983). Functional properties of neurons in the middle temporal visual area of the macaque: I. Selectivity for stimulus direction, speed, and orientation. Journal of Neurophysiology, 49, 1127–1147.

Murphey, D., & Maunsell, J. H. R. (2007, May 15). Behavioral detection of electrical microstimulation in different cortical visual areas. Current Biology, 17(10), 862–867. http://www.ncbi.nlm.nih.gov/entrez/query.fcgi?cmd=Retrieve&db=PubMed&dopt=Citation&list_uids=17462895

Murphey, D., & Maunsell, J. H. R. (2008, May 20). Electrical microstimulation thresholds for behavioral detection and saccades in monkey frontal eye fields. Proceedings of the National Academy of Science USA, 105(20), 7315–7320. http://www.ncbi.nlm.nih.gov/entrez/query.fcgi?cmd=Retrieve&db=PubMed&dopt=Citation&list_uids=18477698

Newsome, W. T., & Pare, E. B. (1988). A selective impairment of motion processing following lesions of the middle temporal visual area (MT). Journal of Neuroscience, 8, 2201–2211.

Ni, A. M., & Maunsell, J. H. R. (2010, May 11). Microstimulation reveals limits in detecting different signals from a local cortical region. Current Biology, 20(9), 824–828. https://doi.org/10.1016/j.cub.2010.02.065

Ni, A. M., Ray, S., & Maunsell, J. H. R. (2012, Feb 23). Tuned normalization explains the size of attention modulations. Neuron, 73(4), 803–813. https://doi.org/10.1016/j.neuron.2012.01.006

Nielson, H. C., Knight, J. M., & Porter, P. B. (1962, Apr). Subcortical conditioning, generalization, and transfer. J Comp Physiol Psychol, 55, 168–173. https://doi.org/10.1037/h0043129

Pasternak, T., & Merigan, W. (1994, May-Jun). Motion perception following lesions of the superior temporal sulcus in the monkey. Cereb Cortex, 4(3), 247–259. http://www.ncbi.nlm.nih.gov/entrez/query.fcgi?cmd=Retrieve&db=PubMed&dopt=Citation&list_uids=8075530

Pi, H. J., Hangya, B., Kvitsiani, D., Sanders, J. I., Huang, Z. J., & Kepecs, A. (2013, Nov 28). Cortical interneurons that specialize in disinhibitory control. Nature, 503(7477), 521–524. https://doi.org/10.1038/nature12676

Popa, L. S., Streng, M. L., Hewitt, A. L., & Ebner, T. J. (2016, Apr). The Errors of Our Ways: Understanding Error Representations in Cerebellar-Dependent Motor Learning. Cerebellum, 15(2), 93–103. https://doi.org/10.1007/s12311-015-0685-5

Rosa, M. G., & Elston, G. N. (1998, Apr 20). Visuotopic organisation and neuronal response selectivity for direction of motion in visual areas of the caudal temporal lobe of the marmoset monkey (Callithrix jacchus): middle temporal area, middle temporal crescent, and surrounding cortex. J Comp Neurol, 393(4), 505–527. https://www.ncbi.nlm.nih.gov/pubmed/9550155

Rust, N., Mante, V., Simoncelli, E., & Movshon, J. (2006, Nov). How MT cells analyze the motion of visual patterns. Nature Neuroscience, 9(11), 1421–1431. http://www.ncbi.nlm.nih.gov/entrez/query.fcgi?cmd=Retrieve&db=PubMed&dopt=Citation&list_uids=17041595

Sadras, N., Pesaran, B., & Shanechi, M. M. (2019, Oct 25). A point-process matched filter for event detection and decoding from population spike trains. J Neural Eng, 16(6), 066016. https://doi.org/10.1088/1741-2552/ab3dbc

Salzman, C. D., Murasugi, C. M., Britten, K. H., & Newsome, W. T. (1992). Microstimulation in visual area MT: Effects on direction discrimination performance. Journal of Neuroscience, 12, 2331–2355.

Tehovnik, E. J., & Slocum, W. M. (2006). Phosphene induction by microstimulation in V1. Brain Research Reviews.

Tootell, R. B. H., Reppas, J. B., Dale, A. M., Look, R. B., Sereno, M. I., Malach, R., Brady, T. J., & Rosen, B. R. (1995). Visual motion aftereffect in human cortical area MT revealed by functional magnetic resonance imaging. Nature, 375, 139–141.

Tootell, R. B. H., Reppas, J. B., Kwong, K. K., Malach, R., Born, R. T., Brady, T. J., Rosen, B. R., & Belliveau, J. W. (1995). Functional analysis of human MT and related visual cortical areas using magnetic resonance imaging. Journal of Neuroscience, 15, 3215–3230.

Watamaniuk, S. N. (1993). Ideal observer for discrimination of the global direction of dynamic random-dot stimuli. J Opt Soc Am A, 10, 16–28. https://doi.org/10.1364/josaa.10.000016

